# Risk of disease spillover from dogs to wild carnivores in Kanha Tiger Reserve, India

**DOI:** 10.1101/360271

**Authors:** V. Chaudhary, N. Rajput, A. B. Shrivastav, D. W. Tonkyn

**Affiliations:** Department of Biological Sciences, Clemson University, Clemson, SC, 29634, USA; School of Wildlife Forensic and Health, South Civil Lines, Jabalpur, M. P., India; Department of Wildlife Ecology and Conservation, University of Florida, Gainesville, FL, 32603 USA; Department of Biology, University of Arkansas at Little Rock, Little Rock, AR 72204, USA

**Author notes:** **Correspondence** V. Chaudhary, Newins-Ziegler Hall,1745 McCarty Road, Department of Wildlife Ecology and Conservation, University of Florida, Gainesville, FL, 32611.

**Keywords:** Canine Distemper Virus, Canine Parvovirus, Canine Adeno Virus, Rabies, Tiger, Dog, India, Wildlife disease

## Abstract

Many mammalian carnivore species have been reduced to small, isolated populations by habitat destruction, fragmentation, poaching, and human conflict. Their limited genetic variability and increased exposure to domestic animals such as dogs place them at risk of further losses from infectious diseases. In India, domestic and feral dogs are associated with villages in and around protected areas, and may serve as reservoirs of pathogens to the carnivores within. India’s Kanha Tiger Reserve (KTR) is home to a number of threatened and endangered mammalian carnivores including tiger (*Panthera tigris*), leopard (*Panthera pardus*), wolf (*Canis lupus*), and dhole (*Cuon alpinus*). It also has more than 150 villages with associated dog populations. We found that dog populations ranged from 14 to 45/village (3.7 to 23.7/km^2^), and did not vary with village area, human population size, or distance from the KTR’s core area, though they all increased between summer 2014 and winter 2015, primarily through reproduction. No dog tested positive for rabies but seroprevalence levels to three other generalist viral pathogens were high in summer (N=67) and decreased somewhat by winter (N=168): canine parvovirus (83.6% to 68.4%), canine distemper virus (50.7% to 30.4%) and canine adenovirus (41.8% to 30.9%). The declines in seroprevalence were primarily due to new recruitments by birth and these were not yet exposed to the viruses. Wild carnivores frequently entered the villages, as shown by tracks, scats, kills and other indicators, and the dogs are known to leave the villages so that encounters between dogs and wild carnivores may be common. We conclude that there is a large population of unvaccinated dogs in and around Kanha Tiger Reserve, with high levels of seroprevalence to pathogens with broad host ranges and these dogs which interact with wild carnivores, therefore posing a high risk of disease spillover to the wild carnivores.

## Introduction

Mammalian wild carnivores, henceforth ‘carnivores’, are threatened by habitat destruction and fragmentation, poaching, and prey depletion (Di Marco *et al.*, 2014). Many persist in small isolated populations, which makes them increasingly vulnerable to infectious diseases caused by pathogens with broad geographic and host ranges (Young, 1994, Thorne & Williams, 1988; Altizer *et al.*, 2003). Indeed, carnivores are threatened by infectious diseases to a greater degree than other mammalian taxa (Pedersen *et al.*, 2007) and, while their typically small populations may not sustain many pathogen species (Lafferty & Gerber, 2002), they are vulnerable to disease spillover from domestic carnivores such as dogs (*Canis familiaris*).

Dogs have extensive geographical ranges and large populations and serve as reservoirs for pathogens including rabies, canine parvovirus (CPV), canine distemper virus (CDV) and canine adeno virus (CAV) that can infect a wide range of carnivores (*e.g*., SilleroZubiri, King & MacDonald, 1996; Cleaveland *et al.*, 1999, 2000, 2001; Alexander *et al.*, 2010; Gompper, 2014). Transmission from dogs to carnivores can occur directly or indirectly, through mating (Bohling & Waits, 2011), predation, or co-scavenging (Butler, Du Toit & Bingham, 2004; Fiorello *et al.*, 2006; Young *et al.*, 2011), and has been implicated in wildlife disease outbreaks around the world (Table 1). Therefore, knowledge of disease prevalence in dogs near wildlife protected areas is essential for conservation planning. Such information is sparse from densely populated countries like India, which has fourth largest dog population in the world (Gommper 2014), lot of which are unvaccinated and live close to wildlife protected areas.

**Table 1.**
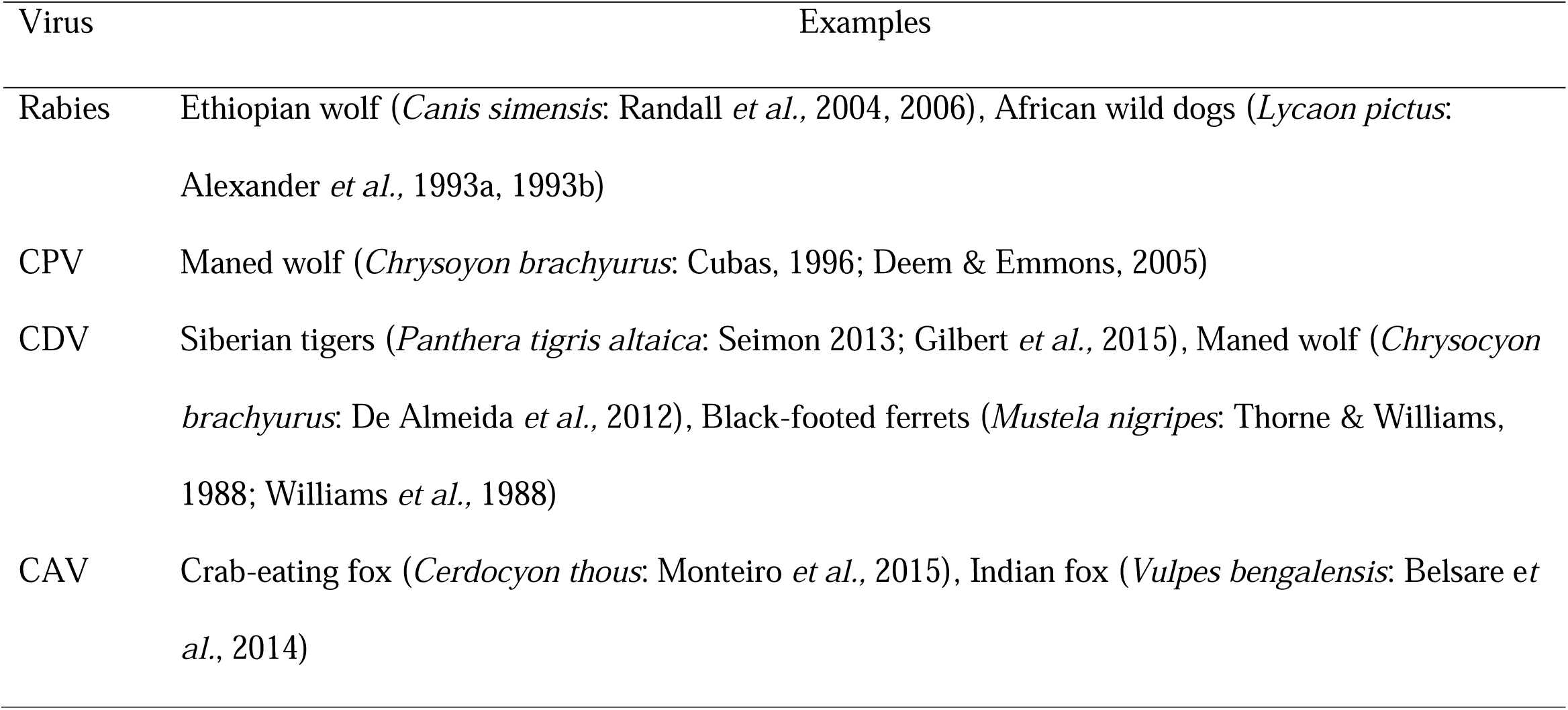
Examples of viral infections in wild carnivores that were contracted from domestic dogs.

| Virus | Examples |
| --- | --- |
| Rabies | Ethiopian wolf (*Canis simensis*: Randall *et al.*, 2004, 2006), African wild dogs (*Lycaon pictus*: Alexander *et al.*, 1993a, 1993b) |
| CPV | Maned wolf (*Chrysoyon brachyurus*: Cubas, 1996; Deem & Emmons, 2005) |
| CDV | Siberian tigers (*Panthera tigris altaica*: Seimon 2013; Gilbert *et al.*, 2015), Maned wolf (*Chrysocyon brachyurus*: De Almeida *et al.*, 2012), Black-footed ferrets (*Mustela nigripes*: Thorne & Williams, 1988; Williams *et al.*, 1988) |
| CAV | Crab-eating fox (*Cerdocyon thous*: Monteiro *et al.*, 2015), Indian fox (*Vulpes bengalensis*: Belsare *et al.*, 2014) |

In this study we assess the threat of rabies, CPV, CDV, and CAV spillover from dogs to carnivores in a tiger (*Panthera tigris*) conservation priority area in central India - Kanha Tiger Reserve (KTR). KTR supports a number of threatened mammalian carnivores including tiger, leopard (*Panthera pardus*), wolf (*Canis lupus*), and dhole (*Cuon alpinus*). It also contains approximately 150 villages in the buffer zone (1,134 km^2^), where regulated human activities are allowed, and eight villages in the core zone (‘core’) (940 km^2^), where stricter regulations on human activity are implemented. All these villages support dog populations (DeFries, Karanth & Pareeth, 2010). In India, dogs near protected areas threaten wild animals through predation and competition (Home, Bhatnagar & Vanak, 2017); and may serve as reservoirs of pathogens that threaten wild carnivores. Majority of dogs in KTR are semi-owned and associate with household but are unrestrained. A few are owned and restrained, and the rest are feral. No village dogs were vaccinated (V. C. pers. comm. with park authorities and owners) so they pose a potential disease spillover risk to carnivores.

Carnivores in KTR frequent the villages to prey on dogs (Karanth *et al.*, 2013) and livestock (Miller *et al.*, 2015), while dogs enter the core with or without their owners. As a result, these animals can interact directly through scavenging on carcasses or predation or, indirectly through scats or spray marks; in ways that can transmit pathogens. We predict that these interactions will be more frequent in villages that are closer to the core (as in Chile) (Torres & Prado, 2010) or that are larger since they may support more dogs. Finally, as some pathogens have common routes of transmission or suppress the immune response, we predict that there will be elevated rates of co-exposure, amplifying the threat (Griffiths *et al.*, 2011). We conducted this study in three parts over two field periods. First, we estimated the abundance of dogs and compared their numbers in total and by sex and age in villages of varying sizes, season and, distances of the village from the core. Second, we measured the exposure of dogs to rabies, CPV, CDV and CAV using seroprevalence of antibodies and, in seropositive cases, by PCR; we compared the prevalence by sex and age of dogs, season and, distances from the core. Finally, we opportunistically recorded the occurrence of wild carnivores in the villages as a proxy for contact rate between dogs and carnivores. We analyzed the results to estimate the temporal and spatial variation in the threat of disease transmission from dogs to carnivores.

## Materials and methods

### Study area

KTR (22^0^7‘N– 22^0^27‘N, 80^0^26 ‘E – 81^0^3’E) (Fig. 1) was established as a National Park in 1955 and declared a tiger reserve in 1973 (Damodaran, 2009). It ranges in elevation from 600-870 m a.s.l., has mean monthly temperatures range from 17 ^0^C in winter (October-January) to 32.5 ^0^C in summer (April-June), and receives an average annual precipitation of 1800 mm. KTR’s vegetation is composed of dry deciduous forest (51%), moist deciduous forest (27%), and former agricultural fields now maintained as grasslands. The dry deciduous forest is characterized by the trees *Angoiessus latifolia, Gardenia latifolia, Buchanania lanzan* and *Sterulica urens*, while the moist deciduous forest is dominated by *Shorea robusta, Tectonia grandis*, and *Terminalia tomentos*. The bamboo *Dendrocalamus strictus* dominates the understory in both forests (Newton, 1988). The primary economic activity in the area is cotton and rice farming, small scale livestock management, and wildlife tourism. Farmers suffer economic losses through crop raids by wild herbivores and livestock depredation by carnivores, mainly leopard, tiger, jackal, and dhole (Karanth *et al.*, 2013). Wildlife tourism is regulated by the Forest Department and allowed in the core zone, but other human activities like logging and hunting are prohibited.

**Fig. 1.**
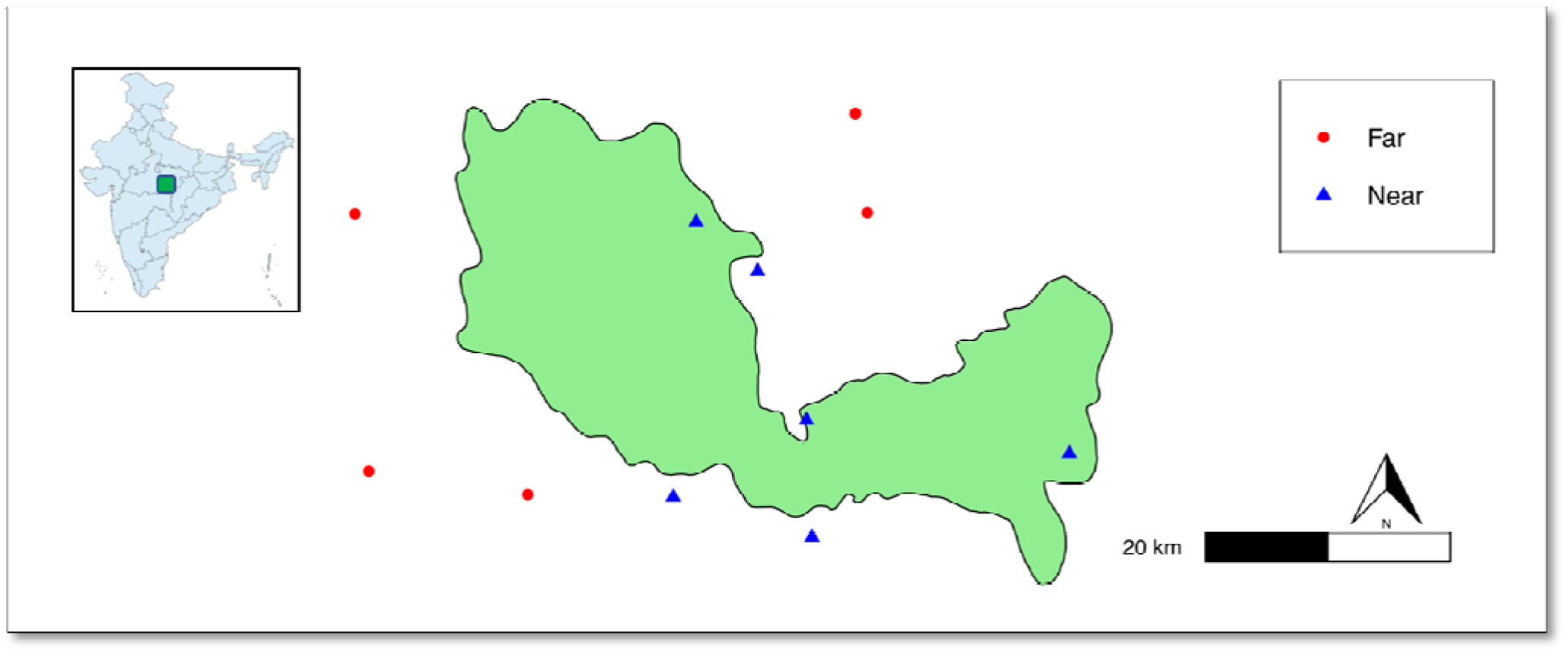
Map of Kanha Tiger Reserve core zone with location of far (>6 km) and near (< 2km) villages that were sampled in summer and winter; the inset shows the location of KTR in India.

### Field surveys

We surveyed dog populations in villages over two field seasons, from May to July 2014 (’summer’) and from January to March 2015 (‘winter’) for abundance, seroprevalence and contact rate with wild carnivores. We randomly selected five villages with centers < 2 km from the core boundary (henceforth ‘near’) and four villages (summer), and then five (winter) with centers > 6 km from the core boundary (‘far’) (Fig. 1). All villages were sampled in both seasons except one near village that was replaced in winter and one far village that was added in winter (Table 2). After establishing the active periods for dogs (6:00-9:00 AM & 5:00-6:30 PM in summer; 7:00-9:00 AM & 4:00-5:30 PM in winter), we conducted non-invasive photographic surveys of dogs using a Nikon D3000 digital camera and 80-200 mm lens, on a motorcycle at ≤ 20 km/h on roads and on foot in alleys. We noted individual dog’s sex (males by descended testicles), age category (juveniles < 1 year or adults by asking the owners and/or estimation), and coat patterns. Each survey was conducted over two consecutive days (a “mark” and “recapture” in each village) in summer and three consecutive days (a “mark” and two morning and two afternoon “recaptures”) in winter. We measured the area of each village by tracking its outermost boundary with a Garmin GPS (Montana 650, Olathe, Kansas, USA). We estimated the dog density by dividing the estimated population size by the village area. The human population size of each village was taken from the 2011 India census (http://censusindia.gov.in/).

**Table 2.** Details of the villages near Kanha Tiger Reserve that were studied, with their human and estimated dog densities which were estimated using program CAPTURE (White & Burnham 1999).

| Village | Distance from KTR<br>core (km) | Area (km<sup>2</sup>) | # humans/km<sup>2</sup> in<br>2011 | Estimated # dogs/km<sup>2</sup><br>Summer 2014 | Estimated # dogs/km<sup>2</sup><br>Winter 2015 |
| --- | --- | --- | --- | --- | --- |
| Bhilwani (N1) | 0 | 3.69 | 141 | 11.6 | 17.4 |
| Lagma (N2) | 1 | 2.06 | 273 | 3.7 | 5.4 |
| Samnapur (N3) | 0 | 3.86 | 253 | 5.1 | 7.7 |
| Lapti (N4) | 0 | 3.17 | 64 | 9.5 | 12.6 |
| Arandi (N5) | 2 | 4.08 | 56 | 9.7 | NA |
| Ranwahi (N6) | 0 | 4.08 | 98 | NA | 19.1 |
| Rajma (F1) | 6 | 5.5 | 205 | 5.6 | 8.2 |
| Parasmau (F2) | 6 | 9.8 | 61 | 2.1 | 2.8 |
| Chartola (F3) | 7.2 | 4.08 | 147 | 17.1 | 23.7 |
| Ghana (F4) | 6.2 | 2.25 | 129 | 11.4 | 14.3 |
| Chiraidongri (F5) | 9 | 4.48 | 192 | NA | 23.1 |

We collected blood samples from dogs in all surveyed villages to estimate the seroprevalence of rabies, CPV, CDV, and CAV. In summer, we collected blood opportunistically from 67 dogs (42 males and 25 females). In winter, we collected blood from five male and five female adults and four male and four female juveniles from each village, except for three near and two far villages where we could only capture three males and three female juveniles. 35 dogs were sampled in both seasons. Dogs less than four months of age were excluded to rule out presence of maternal antibodies (Greene, 1994). Feral dogs were wary of humans and were therefore excluded from the study. Each dog was gently held while a veterinarian collected 4 ml blood from the saphenous vein and transferred it to a 4 ml vacuette tube kept on ice. Samples were transferred to the School of Wildlife Forensic and Health, Jabalpur, India within 24 hours and stored overnight at 4 ^0^C. Serum was separated by centrifugation (CM24, REMI cooling centrifuge, Goregaon E, Mumbai 400063, India) at 3,000g for 15 minutes and stored at −40 ^0^C until further analysis.

### Antibody detection

We tested the sera of all 67 animals collected in summer and one vaccinated dog (positive control) for rabies antibodies with Bio-Rad’s Platelia II test kit (Bio-Rad, Hercules, California, USA). This immune-enzymatic kit uses solid phase inactivated rabies glycoprotein G. Since no sample tested positive except for the control, we did not repeat this test in the winter. We tested all samples from both seasons for antibodies to CPV, CDV and CAV, using BioGal’s Immunocomb canine vaccichek solid phase immunoassay kit (Bio Galed lab, Kibbutz Galed, Israel, 1924000) (Belsare & Gompper, 2013; Waner *et al.*, 2003). It semi-quantifies antibodies and results can be measured by color-comparison provided in the kit. The results are in the form of ‘S’ units from S0 to S6, where S0 indicates no antibodies were detected; S1 and S2 suggest that antibodies were detected in low titers; and S3 to S6 suggest that antibodies are present with minimum 1:16 titer of virus neutralization for CAV, 1:80 titers by hemagglutination inhibition test for CPV, and 1:32 virus neutralization test for CDV. We categorized the results as: seronegative (S0) and seropositive (S1and above; specifically, S1 & S2 = ‘low titers’, S3 = ‘medium titers’ and S4+ = ‘high titers’). All samples seropositive for CPV and CDV were tested for the viral nucleic acid using PCR, along with positive controls. We did not test samples seropositive for CAV for the virus as we lacked a positive control. The methods for DNA and RNA extraction and PCR analyses are described in Supporting Information S1.

### Carnivore encounters

We opportunistically recorded signs of carnivores (direct sightings, photographs, scats and, footprints) in surveyed villages as ‘encounters’ over 60 day periods in both seasons. Naturalists from KTR confirmed the identifications based on scats. On five occasions, we had permission from the KTR Forest Department to install camera traps (Capture IR, 5 MP camera, Cuddeback, Greenbay, WI, USA, 54115) near carcasses of livestock killed by tigers. We recorded the geographical location of each carnivore encounter and collected only once from each location to avoid duplication. This method provides a minimum estimate of the contact rate, since many carnivores are nocturnal and can enter villages undetected.

### Statistical analysis

To estimate the dog abundance in each village and season we used the program CAPTURE (Otis *et al.*, 1978; White & Burnham, 1999), an extension of MARK (version 8.x) (White, 2008). Since, the dogs are territorial (Pal, 2003) and our surveys were conducted over two or three consecutive days in each village, we assumed the individual populations were closed during survey. Also, since we used non-invasive means to identify dogs, we assumed there was no loss of marks and no change in behavior following the first “capture”. Using CAPTURE, we compared four models that differ in their assumptions of capture probability: that it is constant (M_0_), it varies among individuals (M_h_), with time (M_t_), and with both (M_th_), and we selected the model with lowest AIC values.

We used the estimated numbers of dogs in all analyses except when they were separated by sex and age; in those cases, with the smaller numbers of individuals, we used the observed counts in each category. We used simple linear regression to see if estimated dog densities per village were correlated with human densities or village areas. We used the non-parametric Wilcox rank sum test to test for differences in abundance of total dogs (estimated), female dogs (observed), and juvenile dogs (observed) in the following contrasts: a. near vs. far villages in summer, b. near vs. far villages in winter c. summer vs. winter for near villages, and d. summer vs. winter for far villages; where contrast a and b test spatial effect and, c and d test seasonal effect.

We used Fisher’s two-tailed exact test to examine the relation between seroprevalence of each pathogen and sex and age category of the dogs, and also to test non-random patterns of co-exposure. We tested for the difference in seroprevalence for near and far villages (only for winter). We calculated odds ratio (OR) for all significant relationships, which tells the likelihood of seroprevalence given the presence or absence of other conditions, here, the sex and age class of the dogs, and the presence of other pathogens. All analyses were conducted using SAS (SAS Studio 3.4, SAS Institute Inc., Cary, NC, USA).

## Results

The villages surveyed ranged in area from 2.06-9.80 km^2^ and in human population size from 142 to 1,324 (Table 2). We photographed 85 unique dogs in the four near villages and 70 in the four far villages in summer, and 119 and 110 respectively in these same villages the following winter. Model M_0_ had the lowest AIC score for all but two villages in the two seasons (Supporting Information S2), so we used it for all population estimates (Supporting Information S3). These estimates were only slightly higher than the observed values: in summer, ranging from 21-34 in near villages (observed: 17-30) and 19 - 25 in far villages (observed: 14 - 20), while in winter increasing to 28 - 45 in near villages (observed: 24 - 41) and to 27 - 34 in far villages (observed: 24 - 30) (Fig. 2a).

**Fig. 2.**
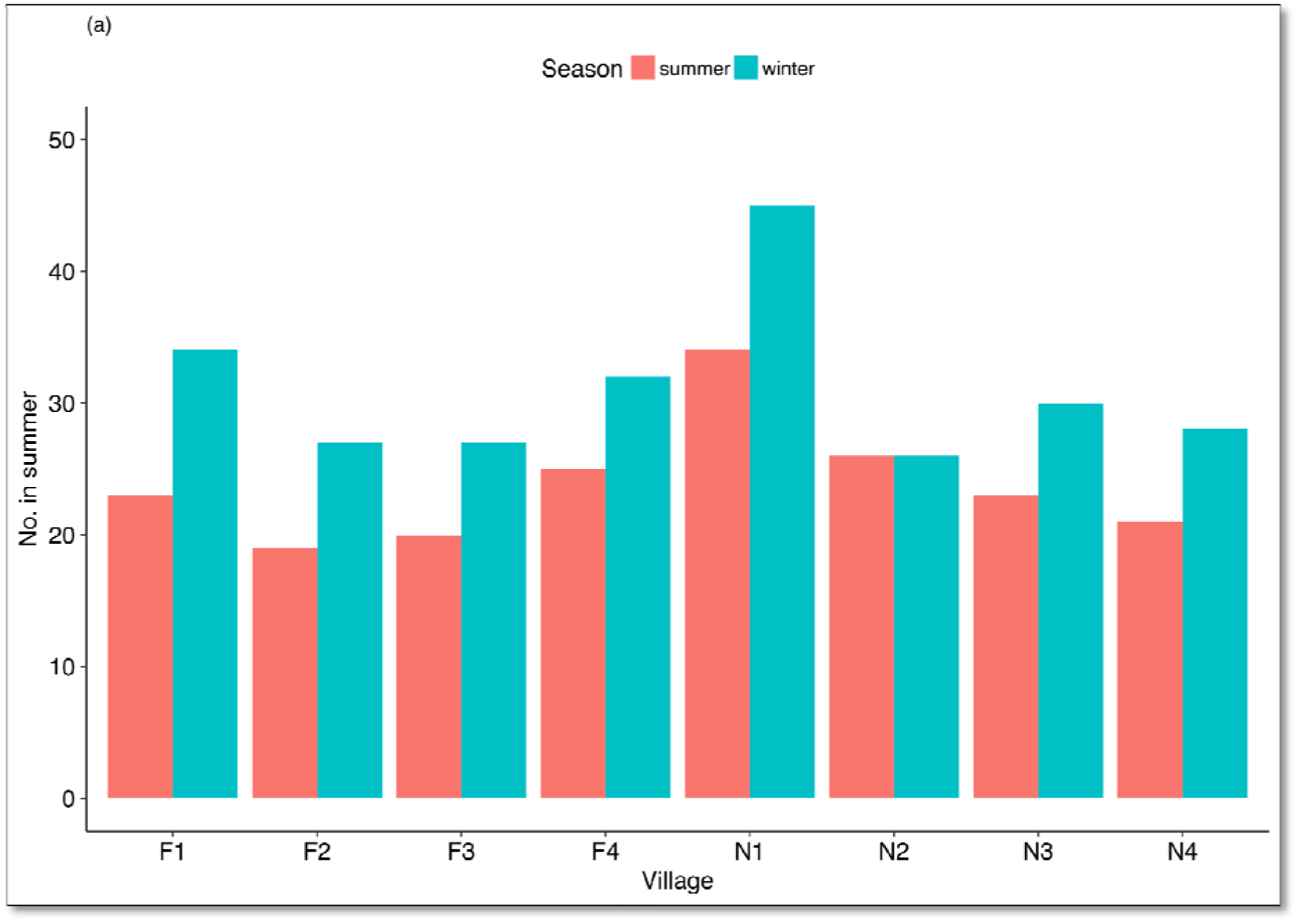
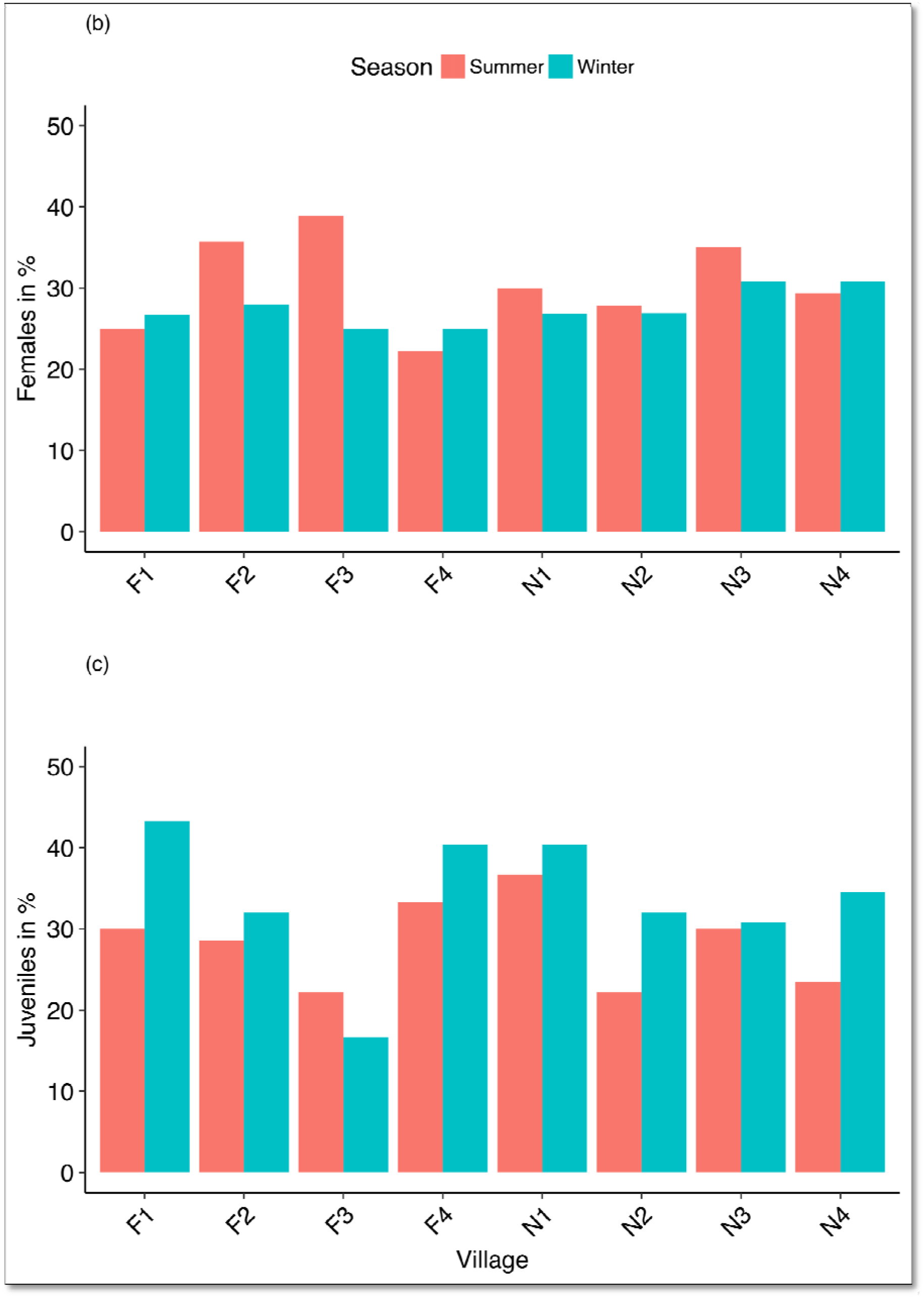
(a) Estimated numbers of dogs in far (F1-F4) and near (N1-N4) villages in summer 2014 and winter 2015. (b) Percent of observed dogs that were females in far and near villages in summer and winter. (c) Percent of observed dogs that were juveniles in far and near villages in summer 2014 and winter 2015.

There was no statistically significant relation between the estimated number of dogs in each village (or the density, accounting for area), and the number or density of humans in each village, or the area of the villages. There was no statistically significant difference in the number or density of dogs in near vs. far villages or in either set of villages in summer vs. winter, though these results are at least partially due to the large variation in the numbers. In fact, the estimated dog density increased from summer to winter in every village surveyed, from an average 10.3/km^2^ in near villages and 12.3/km^2^ in far villages in summer, to 12.2 dogs/km^2^ and 14.3/km^2^ respectively in winter.

Over both field seasons combined, 21% of the dogs were classified as feral, 77% as semi-owned, and 2% as owned. Feral dogs formed 21% and 26% of the dog population in near and far villages respectively in summer, declining to 17% (near villages) and 14 % (far villages) in winter. These differences by proximity to the core and by season were not statistically significant. Females constituted 32.7% and 30% of the population in near and far villages respectively in summer, dropping slightly to 28.5% and 26.2% in winter (Fig. 2b). Juveniles constituted 23% and 28.5% of the population in near and far villages respectively in summer rising to 36.1% and 34% in winter (Fig. 2c). None of these differences, in the number of females or juveniles by proximity to the core or by season, was statistically significant. In the eight villages surveyed in both seasons, 84 of the 155 dogs photographed and identified in summer were still detected in winter (54.2 %). 58% of these 84 dogs were from near villages and 42% were from far villages. The recapture rate was lower for females (32% of 84) and juveniles (40% of 84 recaptures) which were classified as adults in winter.

The seroprevalence of CPV, CDV and CAV were all high in summer and declined in winter: CPV decreased from 83.6% to 68.4%, CDV decreased from 50.7% to 30.4% and CAV decreased from 41.8% to 30.9%. There was no significant effect of age or sex to the seroprevalence in all villages in summer (Table 3) and in near villages in winter (Table 4). However, in far villages in winter, adults were more likely to be seropositive than juveniles for CPV (P = 0.003), CDV (P = 0.003) and CAV (P = 0.01) (Table 4). The proportion of the dogs with antibody titers of S4+, S3 and S1 varied with the pathogen (Supporting Information S4). Dogs that tested positive for CPV antibodies were most often in the high range (S4+) in both seasons, whereas dogs that tested positive for CDV and CAV were predominantly in the moderate range (S3) in both seasons except for dogs in near villages that were more often S4+ in greater proportion as compared to far villages in winter (Supporting Information S5).

**Table 3.** Seroprevalence in various categories of dogs in summer 2014 (S) and winter 2015 (W). This table lists the number of dogs sampled in each season in each category: males, females, adults, juveniles, female adults (FA), female juveniles (FJ), male adults (MA) and male juveniles (MJ), and the percent of each that were seropositive for CPV, CDV and CAV.

| Category | Sample size |  | CPV (%) |  | CDV (%) |  | CAV (%) |  |
| --- | --- | --- | --- | --- | --- | --- | --- | --- |
|  | S | W | S | W | S | W | S | W |
| Total | 67 | 168 | 83.6 | 68.4 | 50.7 | 30.4 | 41.8 | 30.9 |
| Males | 42 | 89 | 83.3 | 62.9 | 52.4 | 29.2 | 40.5 | 33.7 |
| Females | 25 | 79 | 84.0 | 74.7 | 48.0 | 31.6 | 44.0 | 27.8 |
| Adults | 49 | 95 | 87.8 | 77.9 | 46.9 | 40.0 | 46.9 | 43.2 |
| Juveniles | 18 | 73 | 72.2 | 56.2 | 61.1 | 17.8 | 27.8 | 15.1 |
| FA | 18 | 45 | 83.3 | 86.7 | 44.4 | 26.8 | 44.4 | 40.0 |
| FJ | 7 | 34 | 85.7 | 58.8 | 57.1 | 20.2 | 42.9 | 11.8 |
| MA | 31 | 50 | 90.3 | 70.0 | 48.4 | 29.8 | 48.4 | 46.0 |
| MJ | 11 | 39 | 63.6 | 53.8 | 63.6 | 23.2 | 7.1 | 17.9 |

**Table 4.** Seroprevalence in various categories of dogs in near (N) and far villages (F) during winter 2015. This table lists the number of dogs sampled in each season in each category: males, females, adults, juveniles, female adults (FA), female juveniles (FJ), male adults (MA) and male juveniles (MJ), and the percent of each that were seropositive for CPV, CDV and CAV.

| Category | Sample |  | CPV (%) |  | CDV (%) |  | CAV (%) |  |
| --- | --- | --- | --- | --- | --- | --- | --- | --- |
|  | N | F | N | F | N | F | N | F |
| Total | 85 | 83 | 74.1 | 62.6 | 17.7 | 42.9 | 23.5 | 38.6 |
| Males | 46 | 43 | 71.7 | 56.8 | 17.4 | 43.2 | 28.3 | 36.4 |
| Females | 39 | 40 | 76.9 | 69.2 | 18.0 | 43.6 | 18.0 | 41.0 |
| Adults | 50 | 45 | 78.7 | 76.6 | 23.4 | 57.4 | 36.2 | 51.1 |
| Juveniles | 35 | 38 | 68.4 | 44.4<sup>*</sup> | 10.5 | 25.0<sup>*</sup> | 7.9 | 22.2<sup>*</sup> |
| FA | 23 | 22 | 82.6 | 86.4 | 27.1 | 47.1 | 27.1 | 59.1 |
| FJ | 16 | 18 | 68.7 | 47.0 | 18.8 | 23.5 | 18.8 | 17.6 |
| MA | 24 | 25 | 75.0 | 68.0 | 28.2 | 56.0 | 28.2 | 44.0 |
| MJ | 19 | 20 | 68.2 | 42.1 | 25.9 | 26.3 | 25.9 | 26.3 |

\* Statistically significant relation

In winter, dogs that were seropositive for any one pathogen were more likely to be seropositive for one or both of the others (Table 5). This trend was statistically significant in far villages, and near and far villages combined, but not in near villages alone. In the far villages, dogs that were seropositive for any combination of two out of three viruses was more likely to be positive for third one. For example, if a sample tested seropositive for CPV and CDV, it is 13.72 times more likely to be seropositive for CAV (P = 0.001, OR = 13.72).

**Table 5.** Test of relation between seroprevalence for each pathogen (outcome) and seroprevalence of one or both of the other two pathogen (condition) listed for categories: S (summer 2014) total (near and far villages combined), W (winter 2015) total (near and far villages combined), winter 2015 (only near villages) and winter 2015 (only far villages), described using P value, odds ratio and, 95% confidence limits of the odds ratio.

| Pathogen | Condition | P value | OR | 95% confidence limits |
| --- | --- | --- | --- | --- |
| S/total |  |  |  |  |
| CPV | CDV and CAV | 0.304 | 2.29 | 0.52 - 10.07 |
| CDV | CAV and CPV | 0.182 | 2.53 | 0.68 - 9.41 |
| CAV | CDV and CPV | 0.22 | 4.90 | 0.55 - 43.05 |
| W/total |  |  |  |  |
| CPV | CDV and CAV | 0.001<sup>*</sup> | 13.72 | 3.15 - 59.28 |
| CDV | CAV and CPV | 0.001<sup>*</sup> | 4.16 | 1.95 - 8.88 |
| CAV | CDV and CPV | 0.006<sup>*</sup> | 3.12 | 1.34 - 7.25 |
| W/ near |  |  |  |  |
| CPV | CDV and CAV | 0.117 | 4.40 | 0.53 - 36.01 |
| CDV | CAV and CPV | 0.296 | 2.04 | 0.66 - 6.24 |
| CAV | CDV and CPV | 0.769 | 1.38 | 0.40 - 4.73 |
| W/ far |  |  |  |  |
| CPV | CDV and CAV | 0.001<sup>*</sup> | 39.77 | 5.02 - 314.7 |
| CDV | CAV and CPV | 0.001<sup>*</sup> | 9.37 | 3.22 - 27.27 |
| CAV | CDV and CPV | 0.0008<sup>*</sup> | 6.73 | 2.06 - 21.96 |

<sup>\*</sup> Statistically significant relation.

Of the 35 dogs that were surveyed and sampled in both seasons, all 19 dogs that were seropositive for CPV in summer remained so in winter, while 7 more (43.7% of 16) became seropositive by then. Similarly, all 13 dogs that were seropositive for CDV in summer remained so in winter, and three more showed positive seroconversions (13% of 22). For CAV, of nine dogs that were seropositive in summer, one showed negative seroconversion (11.1%) while one of the others showed a positive seroconversion (3.9%) by winter. Results of the RT-PCR tests, indicated active CDV infection for a two-year-old female and a six-year-old male and active CPV infection in an eight-month old male dog (Supporting Information S1).

We had direct and camera trap sightings of wild carnivores within the villages, as well as a variety of indirect records including calls, footprints and scats (Table 6). There were more such records in near villages (16 summer, 18 winter) than in far villages (6 summer, 4 winter).

**Table 6.** Signs of wild carnivores in surveyed villages. Number of signs of wild carnivore presence noted in surveyed near and far villages in summer 2014 (S) and winter 2015 (W).

| Village (type/season) | Type (number) |
| --- | --- |
| Near/S | Animals sighted (2 jackals, 1 leopard, 1 tiger); footprints and drag marks (3 tigers, 1 jackal); scat (6 jackals, 1 leopard), distinctive call (2 jackals) |
| Near/W | Camera trap (1 wild dog, 1 jackal), livestock predation in front of the owner's house (4 tigers), footprints (3 tigers, 1 leopard, 1 jackal), animal sighted (2 tigers), scat (1 tiger, 2 jackals), distinctive call (1 fox, 1 jackal) |
| Far/S | Animal scat (3 jackals), animal sighted (2 jackals), photographic report (1 fox) |
| Far/W | Animal sighted (3 jackals), footprints (1 jackal) |

## Discussion

The potential of disease spillover from dogs to carnivores in KTR is significant as dogs are present in large numbers, are exposed to infectious viruses, and come in direct or indirect contact with carnivores. The average number of dogs in KTR villages that were sampled in both seasons, was 21 in summer and 17 in winter in case of near villages, and in the far villages there were 29 dogs in summer and 27 dogs in winter. The seroprevalence of CPV, CDV, and CAV represents the proportion of the population that were exposed to those viruses at some point and possess antibodies (Greiner & Gardner 2000a, b). While rabies was not detected, the dogs had high seroprevalence to all three viral pathogens in summer, declining slightly in winter with the influx of new juveniles. Finally, there were frequent occasions recorded of direct and indirect contact between carnivores and dogs.

Photographic mark-recapture methods have previously been used to estimate the abundance of dogs and other species with unique coat markings (Lubow & Ransom, 2009; Belsare & Gompper, 2013). Our surveys had a single recapture in summer and four recaptures in winter, and yielded slightly higher estimates than the observed number of dogs in each village, with small standard errors. This indicates that most dogs were observed and identified in the surveys. The densities that we estimated, of 3.7 to 17.1 dogs/km^2^ in summer and 5.4 to 23.7 dogs/km^2^ in winter, are higher than those from villages in the Sikhote-Alin Biosphere Zapovednik in Russia (5/km^2^, Gilbert *et al.*, 2015) but much lower than those from villages of the Great Indian Bustard Sanctuary (GIB) in India (719 dogs/km^2^, Belsare & Gompper, 2013). This may be due to the much larger human population of surveyed villages in GIB supporting dogs (2,973 - 7,448) and the absence of leopards (Belsare & Gompper, 2013) which can prey on dogs (Athreya *et al.*, 2016).

The abundance of dogs per village did not vary with the area, human population size, or distance of the village from the core. The proportions of females and of juveniles, also did not vary significantly with any of these variables. Males were always more abundant than females, suggesting either higher survival or activity rates (Acosta-Jamett *et al.*, 2015). Only 32% of females and 40% of juveniles observed in summer were observed as adults again in winter, suggesting high female and juvenile mortality, though new births elevated the population sizes in all villages surveyed.

Canine rabies is a serious public health concern in India where dog cause 91.5 % of the 15 million animal bites and where 20,000 people die of rabies annually (36% of the world total) (WHO 2016, Knobel et al. 2005). Rabies transmission from dogs therefore likely poses a serious threat to carnivores in India but was not detected in our study because it is highly and quickly lethal (Tepsumethanon *et al.*, 2004). However, most dogs had been exposed to CPV (83.6% seropositive in summer, 68.4% in winter), at rates similar to those reported in GIB (88%: Belsare & Gommper, 2013), Chile (74%: Acosta-Jamett *et al.*, 2015) and Uganda (83%: Millan *et al.*, 2013). These high values reflect the hardiness of CPV, which can survive in the environment for months, and can be transmitted through contaminated soil and arthropodic vectors (Bagshaw *et al.*, 2014). Seroprevalence in juveniles was higher in winter (72% compared to 56.2% in summer) suggesting that it quickly colonizes susceptible hosts, helping ensure its persistence in the population. CPV seroprevalence was higher in the near villages, where carnivores enter frequently (Miller *et al.*, 2015) and may get infected. Of the dogs that tested positive for CPV, the greatest number had S4+ titers suggesting that they have suffered from mild disease with complete recovery. This also suggests that these dogs may have had repeated exposure to the virus and have recovered.

CDV seroprevalence was also high (50.7% in summer, 30.4% in winter) and at rates comparable to those found in Chile (47%: Acosta-Jamett *et al.*, 2015) but lower than those found in GIB (73%: Belsare & Gommper, 2013) and Uganda (100%: Millan *et al.*, 2013). It was also low compared to CPV in KTR. This could either mean that the transmission of the CDV is low in the region or the resulting mortality is high. CDV seroprevalence in near villages in winter (17.7%) was much lower than in far villages (42.9%), making dogs in near villages susceptible to an epidemic of CDV when the conditions for virus become favorable. Proximity of the susceptible host population to carnivores increases the risk of introduced infection to the carnivores. Similar scenario exists for CAV seroprevalence (41.8% in summer and 30.9% in winter), which is low in comparison to GIB (68%: Belsare & Gompper, 2013). CAV is stable in the environment for days, and infected dogs can excrete the virus in urine for at least 6 months (Greene, 1994). Seroprevalence of both CDV and CAV were higher in summer, primarily because of much higher seroprevalence in juveniles in summer than in winter. Despite evidence of exposure of CPV and CAV in KTR dogs, the PCR analyses detected active infection of CPV and CDV in just four animals. It should be noted that seroprevalence can vary with many factors including sex, as pathogens may infect one gender more than the other (Sheridan et al., 2000), interactions in the host community, and existing infections (Glen & Dickman 2005). High seroprevalence may indicate high transmission rates of the pathogen and/or high post-exposure survival rates. To decipher this difference, long term monitoring of the population is required

Pathogens such as CDV have complex relationships with the host and their pathogenesis differs significantly based on the region. CDV infections have resulted in mortality of lions in the Serengeti region (Roelke-Parker *et al.*, 1996), however in southern Africa, lions have existed with CDV without any significant impact despite high exposure (Alexander *et al.*, 2010). One of the factor responsible for the difference is co-infection as the mortality of lions in Serengeti was due to CDV’s association with *Babaesia* (Munson *et al.*, 2008). Co-infection is an important factor in mass die-offs (Goller *et al.*, 2010) and we observed a statistically significant association between the seroprevalence of each of the pathogen with the other two in dogs of far villages in winter. The co-infection can be facilitated through common mode of transmission of pathogens or through immunosuppression (Sykes, 2010) and enablement of secondary infections (Holzman, Conroy & Davidson, 1992). The pathological mechanism of co-exposure or co-infection is beyond the scope of this study. However, the complex disease dynamic of generalist pathogens such as CDV and associated viruses make further research essential for understanding the disease ecology of dogs and carnivores in KTR.

Pathogen spillover from dogs to carnivores in KTR is highly likely as our results confirm the presence of carnivores in the surrounding villages primarily in near villages. These observed contact rates reflect minimum values, as they do not include undetected carnivore signs. Though prohibited, villagers and their dogs enter core zone occasionally and this is true of other protected areas in India as well, where carnivores are surrounded by densely packed human habitations and associated dog populations. Disease transmission in a multi-host system is greater than single host system (Craft *et al.*, 2008) and transmission depends on contact rate, social behavior, and spatial distribution of hosts (Dobson, 2004). This has serious implications for the carnivores of KTR, as dogs here are carriers of pathogens, are largely unrestrained and can infect wild carnivores. Unrestrained dogs, irrespective of the ownership status, form about 75% of the global dog population (WSPA, 2011) and pose threat to wildlife. Feral dogs in particular are bigger threat as they are more prone to suffer from high mortality, malnutrition, and disease (Sowemimo, 2009). Very few dogs photographed in summer were re-sighted in winter in our survey, possibly due to high mortality of these dogs. Our study does not incorporate any seroprevalence data from feral dogs, which may be higher along with the associated mortality. Feral dogs may have more frequent contact with carnivores making them a greater unknown threat.

Fragmentation of carnivore habitats due to anthropogenic activities leads to intense interactions between humans, domestic carnivores and wild carnivores (Thorne & Williams, 1988, Holmes, 1996). These interactions can lead to disease spillover and the risks of infection which can be amplified by environmental stressors like drought, existing infection, prey depletion (Evermann, Roelke & Briggs, 1986) and climate change (Munson *et al.*, 2008). Canine diseases like rabies are public health concern as well (Sudarshan *et al.*, 2007, Sakai *et al.*, 2013). Dogs which form an important link of pathogen transmission between wild carnivores, humans and livestock (MacPherson, 2005) are present in villages of KTR and the disease spillover risk threat caused by these abundant unvaccinated dogs should not be underestimated. Anthropogenic activities have led to decline in species with important ecosystem services such as vultures, which further enhances risk of disease spillover. Vultures rapidly scavenge carcasses thereby limiting the spread of pathogens (DeVault, Rhodes & Shivik, 2003) and decline in vulture population (*Accipitridae* & *Cathridae* family) increases pathogen transmission between carnivores and dogs as overall time spent scavenging is higher and contact between different carnivore species is greater (Ogada & Buij, 2011). This may be true for KTR as well, since vulture sightings there decreased dramatically from initial observations in 2004 until the time of this study (A. B. S. and D.T., pers. observation).

Our results show that dogs in KTR are abundant, are exposed to CPV, CDV and CAV, and come in frequent contact of carnivores. Despite the low densities, these dogs have high turnover adding constant susceptible hosts of the pathogens, maintaining the risk of disease spillover to wild carnivores (Gascoyne *et al.*, 1993; Roelke-Parker *et al.*, 1996). Epidemic events also depend on immunological status and exposure of the pathogen to wild carnivores. We detected exposure of CPV and FPV in one tiger in KTR (Chaudhary *et al.*, in prep), and we recommend continued surveillance of wild carnivore for these pathogens. There is a need of continued investigation of ecology of KTR dogs including their movements patterns and infection recovery rates in KTR, as these estimates are essential to develop disease-dynamic models. Based on our findings we recommend management of infection risks posed by dogs in KTR to prevent pathogen spillover to carnivores. Management options could include culling and vaccination of dogs. Culling though useful to control diseases (Barlow, 1996), can result in compensatory recruitment of dogs and raises ethical concerns in India. Vaccination of reservoir population is an effective management strategy and has resulted in the elimination of diseases like rabies in western Europe and the Serengeti ecosystem (Cliquet 2004; Lembo *et al.*, 2010). Specifically, for population with low pathogen exposure like in KTR (for CDV and CAV), vaccination can be beneficial (Belsare & Gompper, 2015). Combination of vaccination and contraception in dogs reduces disease spread (Carroll *et al.*, 2010) and we recommend the use of oral baiting to deliver both kinds of compounds for disease control in dogs in KTR.

## ACKNOWLEDGMENTS

We thank Clemson University and the National Wildlife Refuge Association for providing funds for the study, the Madhya Pradesh Forest Department particularly Mr. Narendra Kumar and Dr. Suhas Kumar for issuing permits, and the Kanha Tiger Reserve Forest Department particularly Mr. J. S. Chauhan, Dr. Sandeep Agrawal, Dr. Rakesh Shukla and Mr. Rajneesh Singh for providing samples and support. We thank Dr. Kajal Jadhav, Dr. Himanshu Joshi, Dr. K. P. Singh, Dr. Amitod, Dr. Amol Rokade and Dr. Sunil Goyal of the Centre for Wildlife Forensic and Health in Jabalpur, India and Dr. Dharmaveer Shetty of the University of California, Davis, USA for their help in the laboratory and field. We thank Mr. Amit Sankhala and Mr. Tarun Bhati for logistical support.

Surveys and sample collection for this study were conducted in accordance with Institutional Animal Care and Use Committee number 2014-025 of Clemson University and Madhya Pradesh Forest Department’s research permit number I/3023.

